# Low-dimensional prefrontal representations of objects during working memory

**DOI:** 10.64898/2026.06.03.729348

**Authors:** Danai G Sakelliadou, Scott L Brincat, Mikael Lundqvist, Earl K Miller

## Abstract

There is an ongoing debate regarding the dimensionality of neural representations. Some accounts emphasize representation within low-dimensional subspaces or manifolds, while others suggest high-dimensional neural codes where neurons respond independently. Here, we investigate the dimensionality of prefrontal cortex (PFC) representations of visual objects held in working memory. We found that object representations are low-dimensional, occupying only 3–6 effective dimensions during both encoding and maintenance in working memory. Control analyses indicate this dimensionality was not limited by the number of objects tested (40) or neurons sampled (∼100). We also compared PFC dimensionality to that of a well-established deep neural network model of its inferotemporal (IT) inputs and found an approximately 7-fold dimensionality reduction in PFC. These results suggest object representations are compressed into a low-dimensional manifold in PFC, which might be related to attractor dynamics for working memory, and might facilitate interaction with other variables and cognitive control.

**Author summary:** Competing accounts of cortical neural coding propose that neurons either activate in structured, coordinated patterns like schools of fish, or that they activate independently, like hunting sharks. A key metric discriminating these schemes is their effective dimensionality — the number of independent activity patterns needed to characterize information conveyed in a neural population. To test this, we measured the dimensionality of primate prefrontal cortex during working memory for objects, a domain where the inputs are intrinsically complex and high-dimensional. We found that prefrontal dimensionality was surprisingly low, spanning only a handful of dimensions. Using convolutional neural network models of its visual cortex inputs, we found evidence for a substantial prefrontal compression of dimensionality. We hypothesize that this reformatting of high-dimensional sensory information into a compact code can facilitate flexible combination with information from other domains, as proposed by “mixed selectivity” theories of prefrontal coding.

## Introduction

During visual working memory, prefrontal cortex (PFC) must encode, maintain, and manipulate information about objects while supporting flexible decision-making related to them [1]. How object representations are organized at the population level, whether they are held in a high-dimensional or a low-dimensional code, and how their representational geometry changes over time, remains under debate.

Recent work has approached this question by characterizing working-memory activity in terms of manifolds and trajectories within the state space of all possible population spiking activity patterns [2–4]. In recurrent neural network models, working memory representations are often represented by low-dimensional manifolds, such as ring attractors or slow manifolds [5–7]. Consistent with this, many studies have shown that population activity in prefrontal cortex exhibits structured patterns of correlation that organize it into low-dimensional subspaces or manifolds [8–14]. Similar results have been observed in other frontal cognitive and motor-related regions [15–18]. Explicitly or implicitly, these results imply a low-dimensional population representation (Fig 1A, right).

**Fig 1.**
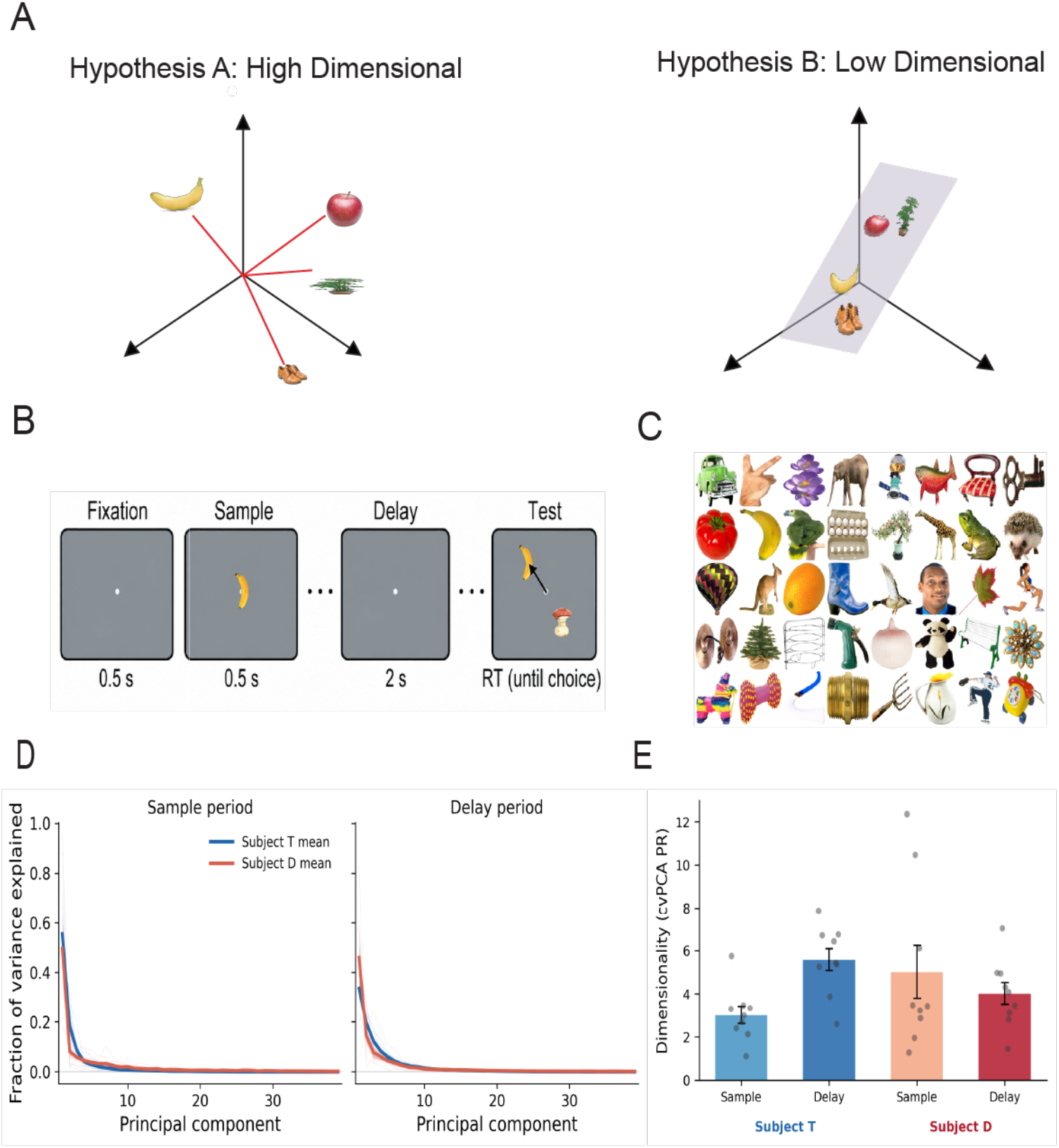
Object representations occupy a low-dimensional space in prefrontal cortex. (**A**) Schematic of the two competing hypotheses. Under a high-dimensional code, each object would be represented along its own independent neural axis, so adding more objects would mean adding more dimensions. In contrast, under a low-dimensional code, different objects are represented as different combinations of a few shared latent dimensions, so many objects can be embedded within the same low-dimensional space. (**B**) Delayed match-to-sample object working memory task. (**C**) Standard set of 40 object images used in the task, consisting of visually distinct real-world objects spanning a broad range of categories, shapes, and colors. Images were licensed from a royalty-free commercial photo database (Hemera Photo-Objects). (**D**) Cross-validated eigenspectra (mean ± SEM across sessions; lighter lines show individual sessions) of PFC activity for the sample and delay epochs. Variance is concentrated in the first few principal components and then falls off quickly for both epochs. **(E)** Cross-validated effective dimensionality (cvPCA-PR) for each subject during the sample and delay epochs. Bars show mean ± SEM across sessions and points show individual sessions. Together, these results indicate that PFC object working memory representations are embedded in a low-dimensional subspace.

In contrast, other studies have proposed that cortical population representations are high-dimensional (Fig 1A, left), with little correlation between the activity of different neurons. High-dimensional codes for sensory stimuli have been observed in lower-level sensory cortical regions [19,20], and it has been suggested that previous results showing low-dimensional representations in prefrontal cortex related to cognitive tasks might be an artifact of limited sampling [20] (but see [21] for an alternative view). Theoretical and empirical work has also suggested prefrontal neurons exhibit “nonlinear mixed selectivity” for different combinations of multiple sensory, cognitive, and motor variables [10,22–25], resulting in a high-dimensional population code [22,26–28]. Thus, there remains a tension between these competing accounts of cortical coding, and the dimensionality of cortical representations is a key measure discriminating between them.

Object working memory in primate prefrontal cortex is a well-studied system [1,29–35] that provides an ideal test case for these questions. The set of all possible real-world objects are a complex, high-dimensional sensory domain [36–40], as are their neural representations in high-level visual areas that provide input to PFC [41]. Thus, it would be unlikely that estimates of population dimensionality are limited by the dimensionality of the stimuli or of upstream sensory processing. Quantifying the dimensionality of PFC object representations could therefore reveal how many dimensions are used to support working memory, whether these dimensions reflect a compression of the upstream object representations, and how the corresponding coding subspace changes between sensory-driven encoding to maintenance. Answers to these questions would provide key constraints on models of working memory.

Here, we address these questions by quantifying the dimensionality of PFC representations for object working memory. We performed large-scale recording of PFC neurons from non-human primates engaged in an object working memory task that sampled a large, diverse set of images of real-world objects. We show that the PFC object representations occupy a relatively low-dimensional space during working memory. The estimated dimensionality was not limited by the number of objects tested or neurons sampled, was substantially lower than a neural network proxy for its IT inputs, and persisted from encoding into the delay despite reorganization of the coding subspace. Together, these results support the hypothesis that PFC maintains object information in a compressed representational format. This low-dimensional object code may facilitate the cross-domain interactions that generate flexible, high-dimensional mixed selectivity codes, reconciling these two seemingly conflicting ideas on neural dimensionality.

## Results

### Object representations occupy a low-dimensional subspace

We recorded from populations of 35–212 neurons (mean 106) in PFC of two non-human primates (NHPs; macaque monkeys) performing a delayed-match-to-sample working memory task (Fig 1B) across 18 sessions (9 sessions for each subject). Each trial, a sample object was shown for 500 ms, and had to be maintained in working memory over a 2 s delay period. Two objects were then displayed, and subjects were required to saccade to the one matching the sample and hold fixation on it for at least one second. The sample and test objects on each trial were selected from a pool of 40 color images of arbitrary real-world objects (Fig 1C). Since dimensionality can be dependent on the complexity of the conditions tested [37], objects were chosen to span a broad range of shapes, textures, colors, and semantic categories, including faces, human and animal bodies, food, flora, tools, vehicles, and other common objects. This same set of objects was used across all sessions, except for a single session that used a partially overlapping set of 40 objects (Supplementary Fig S1).

We aimed to measure the dimensionality of PFC population spiking responses to objects. That is, the dimensionality of the subspace within the space of all possible PFC population activity patterns that specifically codes for objects. Quantifying this using principal components analysis (PCA) on the across-trial mean activity for each object sample condition would contain residual trial-to-trial noise in the estimation of those condition means. Thus, we used cross-validated PCA (cvPCA), a recently developed method that estimates signal dimensionality without contamination from trial-to-trial noise [20]. This yields unbiased principal component eigenvalues by evaluating the between-condition covariance across independent subsets of trials, which have uncorrelated noise (see Methods). Cross-validated eigenspectra revealed that object-related variance was sharply concentrated in the first few dimensions, with subsequent eigenvalues fluctuating around zero (Fig 1D).

We estimated the effective dimensionality of the PFC object representations using the participation ratio (PR) [42]. PR quantifies dimensionality based on the distribution of eigenvalues. It ranges from 1 (when all variance is concentrated in one dimension) to the maximum number of dimensions (when variance is uniformly spread across eigenvalues). In general, the maximum dimensionality is limited by the minimum of the number of conditions tested and neurons sampled. Here, the dimensionality (39 = 40 objects – 1) is limited by the number of objects (except in a single session with 35 sampled neurons). We computed PR on the cross-validated eigenvalues given by cvPCA, constituting an analytical pipeline for computing signal-only dimensionality, which we refer to as cvPCA-PR.

We found that PFC spiking activity had an effective dimensionality of only ∼3–6 dimensions. This was the case during both the sample presentation (subject T: *3*.*01* ± *0*.*40* dimensions, subject D: *5*.*01* ± *1*.*22*; mean±SEM across sessions) and the working memory delay (subject T: *5*.*58* ± *0*.*51*, subject D: *4*.*01* ± *0*.*50*). For both subjects, this was considerably smaller than the maximum possible dimensionality (39 dimensions; *p* < 0.002 for all epochs and subjects, Wilcoxon signed-rank test). It was also significantly greater than the dimensionality measured in data where object condition labels were shuffled across trials, an estimator for the minimal dimensionality given the noise in the data (*p* < *1*.*5* × *10*^−*11*^, Wilcoxon signed-rank test; Fig 1E). These results suggest the PFC neural code for object working memory is low-dimensional, considerably smaller than would be expected if all neurons had independent, uncorrelated responses.

### Dimensionality was not limited by the number of tested conditions

Estimated dimensionality can scale with the number of independent conditions sampled, even beyond the upper bound imposed by the number of conditions [20,37,42]. To determine whether this may have contributed to our finding of low-dimensional PFC coding, we recomputed the effective dimensionality (cvPCA-PR) on randomly selected subsets of 3 to 40 objects. If PFC neurons had independent, uncorrelated selectivity for objects, dimensionality would grow linearly with the number of objects included. This would suggest dimensionality might continue to grow beyond the 40 objects we tested and thus that we may have underestimated PFC dimensionality. If instead objects are embedded in a low-dimensional subspace, dimensionality would eventually saturate with increasing numbers of objects. Distinguishing these scenarios would tell us whether the 3– 6 dimensions we measured truly reflect low-dimensional PFC coding or whether it is a consequence of testing only 40 objects.

For both the sample and delay epochs, effective dimensionality asymptoted well below the 40 objects we tested (Fig 2). Dimensionality initially increased as more objects were added, but then asymptoted around ∼15–30 objects, rather than continuing to grow linearly. To estimate the theoretical dimensionality if an infinite number of objects was tested, we fit this data with saturating exponential curves (all *R*^2^ > 0.97). All four subject/epoch combinations showed clear saturation at 3–6 dimensions, well below the theoretical maximum of 39 (Subject T sample: 3.1[2.4, 4.0] 95% bootstrap confidence interval; Subject T delay: 5.8 [4.7, 6.7]; Subject D sample: 4.4[3.1, 6.1]; Subject D delay: 4.1 [3.2, 4.9]). Thus, our results showing low-dimensional object coding in PFC were not due to the limited number of experimental conditions sampled.

**Fig 2.**
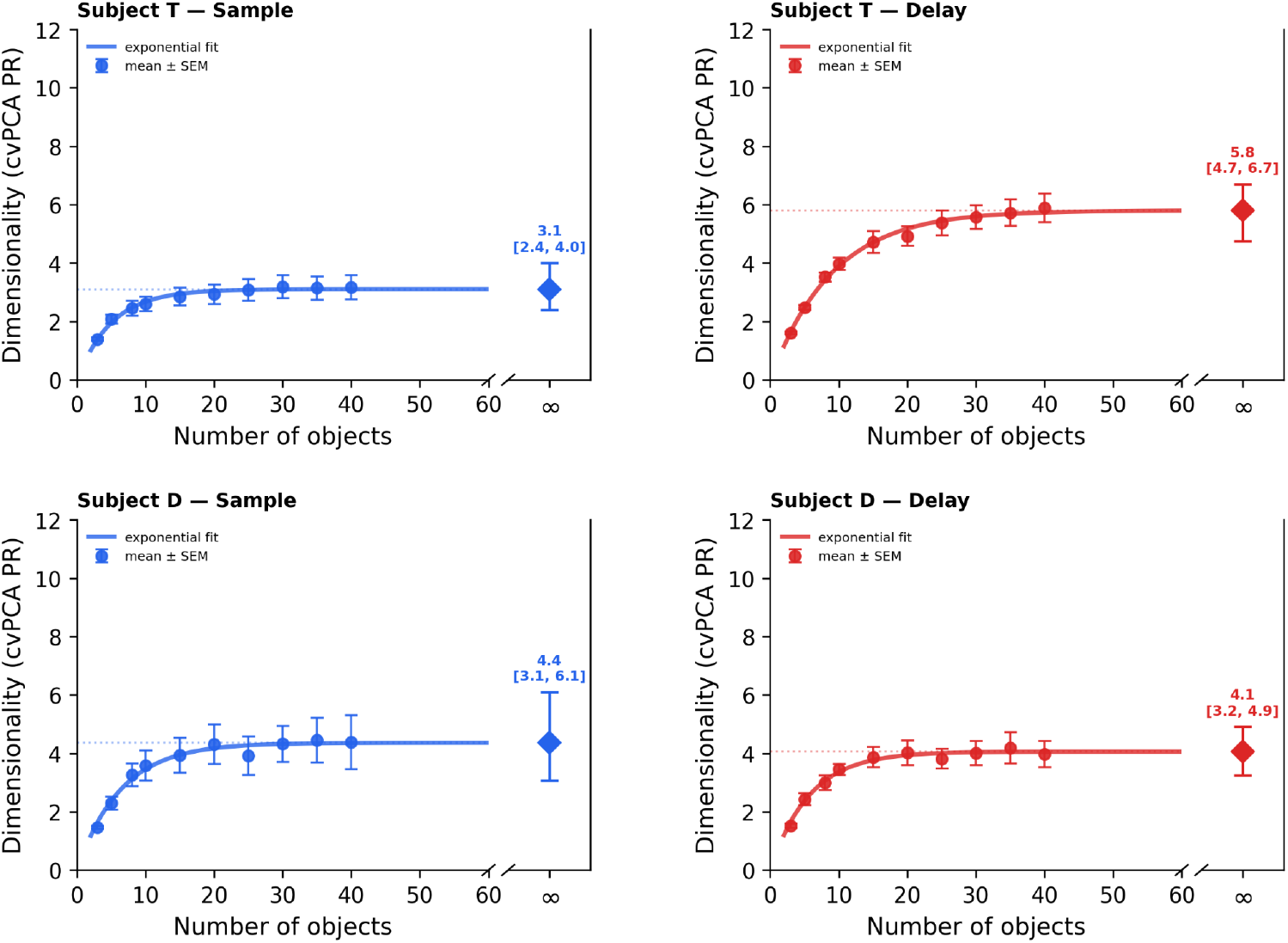
Neural population dimensionality saturates with the number of tested objects. Population dimensionality (cvPCA-PR) as a function of the number of objects tested, for each subject during the sample and delay epochs. Points show the weighted mean ± SEM (weighted by the number of neurons sampled in each session). Lines show saturating exponential fits. Dotted lines and diamonds at right (“∞”) indicate the fitted asymptotic dimensionality and bootstrap 95% confidence intervals. In all cases, dimensionality saturates well below the number of objects tested in the actual experiments (40), indicating this was not a limiting factor in our estimates of dimensionality.

### Dimensionality was not limited by the number of neurons sampled

Estimates of neural population dimensionality can also scale with the number of neurons sampled from the population [20,43,44]. If the intrinsic dimensionality is *D*, we need a number of neurons *N* ≫ *D* to resolve it. Thus, any dimensionality estimate from finite recordings could be limited by the number of neurons rather than reflecting the true population-level value. To determine whether this may have influenced our results, despite sampling ∼100 neurons per session, we randomly subsampled neurons with varying population size and measured effective dimensionality (cvPCA-PR) at each (Fig 3). Because the number of recorded neurons varied across sessions, we limited this analysis to the minimum population size across all sessions for each subject (subject T: 35 neurons, subject D: 118 neurons).

**Fig 3.**
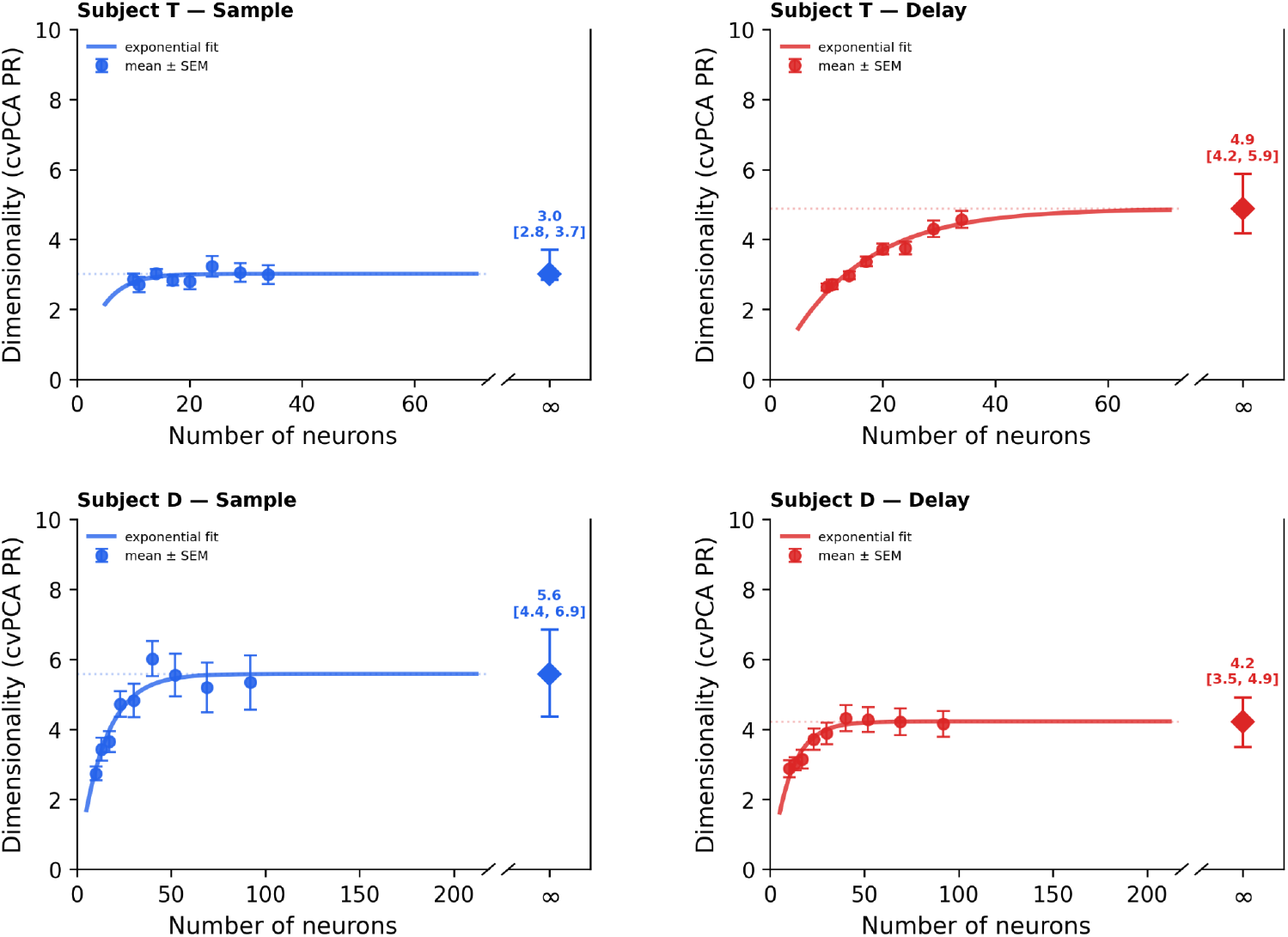
Effective dimensionality saturates with neuronal population size. Dimensionality (cvPCA-PR) of population activity as a function of the number of neurons sampled, for each subject during the sample and delay epochs. Circles indicate weighted mean ± SEM (weighted by the number of sessions contributing to each data point). Lines show saturating exponential fits. Dotted lines and diamonds at right (“∞”) denote the fitted asymptotic dimensionality with bootstrap 95% confidence intervals. In all cases, dimensionality saturates around or below the typical PFC population sizes actually sampled in our data, indicating estimates were not strongly influenced by limited neuronal sampling.

Results suggest that dimensionality had asymptoted within the population sizes we sampled (Fig 3). In most cases, dimensionality initially increased with population size, but then asymptoted around ∼50 neurons, in the range of our smallest PFC sample and much less than our typical sample size. The lone exception was the sample epoch of subject T, where dimensionality was relatively invariant with population size (Fig 3, upper-left). We again fit saturating exponential functions to dimensionality as a function of population size (all *R*^2^ > 0.91, except for subject T sample data: *R*^2^ = 0.28). Results suggest that estimated dimensionality again saturated at around 3–6 dimensions (Subject T sample: 3.0 [2.8, 3.7]; Subject T delay: 4.9 [4.2, 5.9]; Subject D sample: 5.5 [4.4, 6.9]; Subject D delay: 4.2 [3.5, 4.9]). This indicates that our estimates of dimensionality were not limited by the sampled population size.

### Dimensionality remains low-dimensional in larger populations of synthetic neurons

To further confirm that our sampled neural populations are large enough to veridically measure dimensionality, we used a generative model to extrapolate to populations of thousands of simulated neurons. To build a generative model based on the statistical properties of our real data, we first selected high signal-to-noise data to fit the model on. We pooled neurons across all sessions into a nonsimultaneously recorded “pseudopopulation”. For this we used data from subject T, who had overall higher population object decoding accuracy. We then used an iterative pruning procedure to select the neurons most informative about object identity (see Methods). This resulted in a pseudopopulation of *N*=139 informative neurons. We estimated the empirical between-object covariance matrix in this population and simulated synthetic neurons by sampling condition-mean spike rates from a multivariate Gaussian with the same covariance structure. Finally, per-trial spike rates were simulated with a zero-inflated Poisson (ZIP) model [45] fit to the real population’s spike rate statistics. This procedure allowed us to generate a simulated population with the same expected dimensionality as our real data, but with much larger sampled population sizes.

The generatively simulated population had a similar eigenspectrum (Fig 4A) and object decoding accuracy to the empirical population used to generate it. The effective dimensionality of the simulated and real populations was also similar within the range of the empirically sampled population (Fig 4B). Critically, the dimensionality of the generatively simulated population also asymptoted at a similar level, and did not increase, even up to a population size of 3,000 simulated neurons (Fig 4B). These results suggest that prefrontal dimensionality would remain limited to a similar small number of dimensions, even if the entire population was sampled.

**Fig 4.**
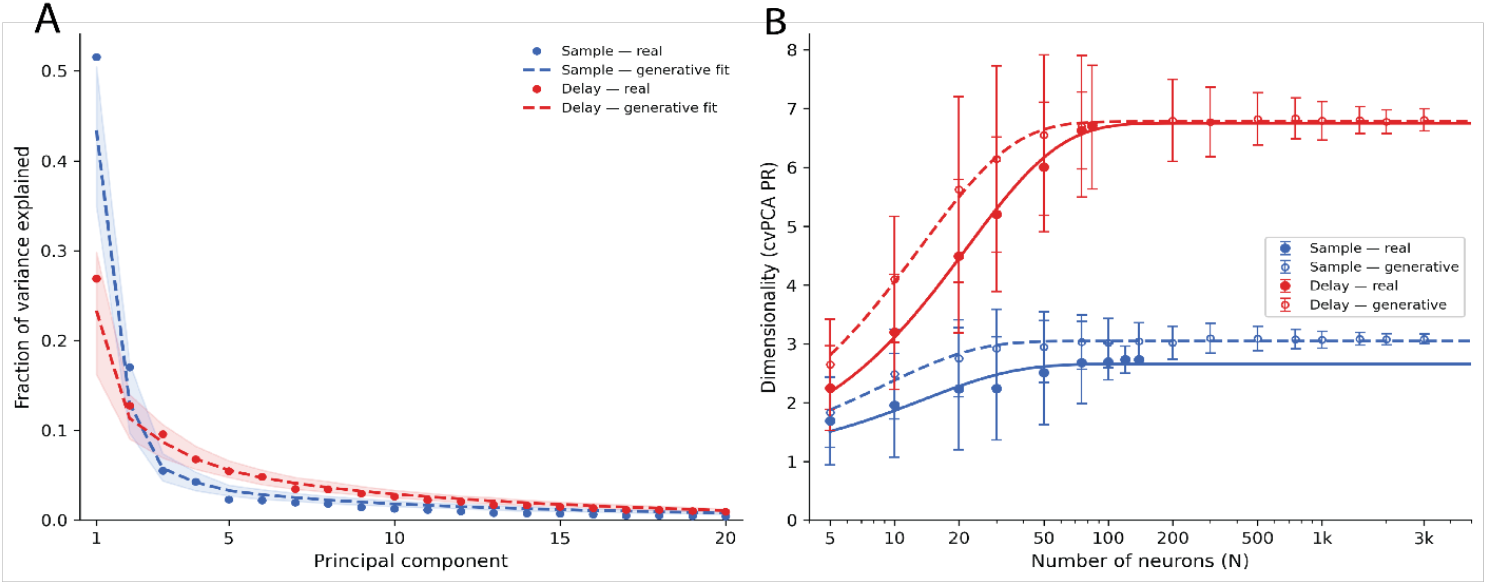
Generative model validation confirms bounded dimensionality beyond the recorded population. (**A**) Fraction of variance explained across principal components, comparing real data (dots) and synthetic populations generated by the fitted generative model (dashed lines: mean, shaded band: 95% percentile interval across 200 simulated populations) during the sample (blue) and delay (red) epochs. In both epochs, the generative model closely reproduces the eigenspectrum of the real data with variance concentrated in the first few components and then decaying rapidly. (**B**) Dimensionality (cvPCA-PR) as a function of population size (N) for Subject T during sample and delay epochs, comparing real vs generative data. Solid curves and symbols show estimates from the real pseudopopulation (note that these values are not the same as those for the full per-session populations in Fig 3; see Methods for details). Dashed lines and lighter symbols show synthetic populations extrapolated to much larger population sizes. Together these results indicate that low dimensionality of object representations is intrinsic and not an artifact of limited recording yield.

To confirm that our cvPCA-PR analysis pipeline is able to capture higher dimensionality when it is present, we also generated synthetic high-dimensional neuronal populations. We used the same generative model as above, but ablated (set to zero) all off-diagonal entries in the estimated between-condition covariance matrix. This manipulation preserves the within-condition variances, but removes the covariance structure that reduces population dimensionality. This resulted in a flattening of the cvPCA eigenspectrum (Fig S2A) and near-maximal estimated cvPCA-PR dimensionality (sample epoch: 36.3; delay epoch: 37.9; Fig S2B). Even in synthesized populations of similar size to our recorded PFC population (∼100 neurons), dimensionality was substantially higher than the real data (∼20–25 dimensions). These results suggest that the low dimensionality we observed in the real PFC data was not due to an inability of our analysis to capture true high-dimensional structure.

### Functional dimensionality is largely consistent with spectral dimensionality

The participation ratio summarizes how variance is distributed across dimensions, but it does not explicitly tell us whether those dimensions carry the information a downstream area could use to read out object information. We therefore confirmed our dimensionality measurements using an alternative method based on linear classification.

We trained linear support-vector machine (SVM) classifiers to decode sample object identity from neural data projected onto its top principal components, testing from 1 to 39 components (Fig 5). Within each cross-validation fold, PCA was computed on condition-averaged activity in the training set. Classification accuracy initially increased with the number of dimensions, then plateaued at around 10 dimensions. We quantified the functional dimensionality as the smallest number of dimensions needed to reach 95% of peak accuracy (max accuracy for subject T: 12.2 ± 0.8% during sample and 13.2 ± 1.3% during delay and subject D:6.6 ± 0.6% during sample and 6.7 ± 0.7% during delay; chance=2.5%). This resulted in functional dimensionality estimates of 6–9 dimensions. The modest gap between functional (∼6–*9*) and spectral (cvPCA-PR; ∼3–*6*) dimensionality may reflect the lack of noise correction in the former, or it may reflect the different ways they quantify dimensionality. Spectral dimensionality reflects how variance is distributed across the eigenspectrum, whereas functional dimensionality reflects all dimensions conveying significant information, so even dimensions carrying small amounts of discriminative signal can result in higher values. Nevertheless, both methods converge on a small number of dimensions in the prefrontal object representation.

**Fig 5.**
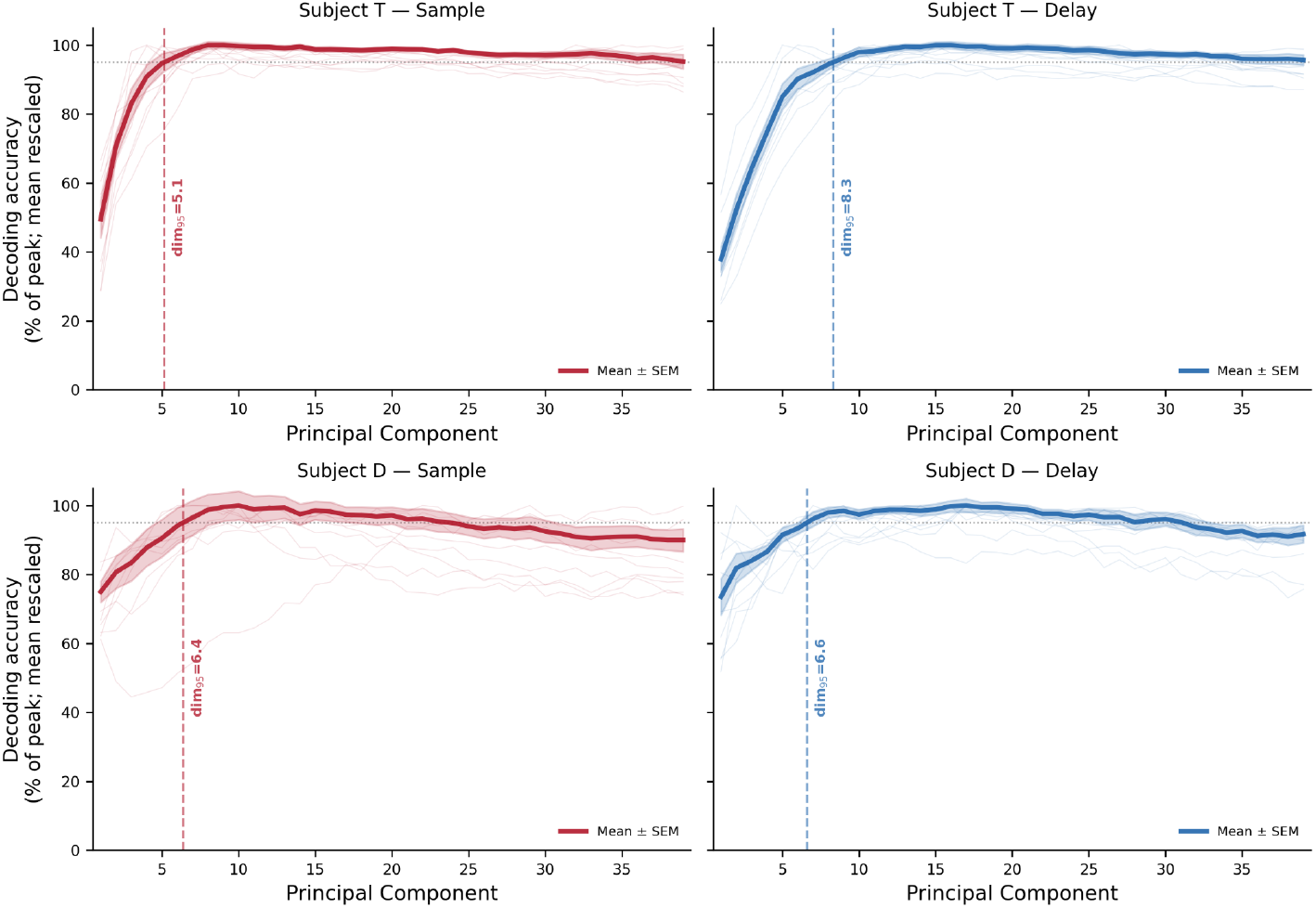
Object decoding accuracy saturates after a small number of principal components. Population decoding accuracy for object identity (40-way linear SVM) is plotted as a function of the number of PCA dimensions retained, shown separately for each subject and task epoch. Faint traces show individual sessions, normalized to their peak. Curves show across-session mean ± SEM of normalized accuracy. Vertical dashed lines indicate the functional dimensionality, defined as the smallest number of dimensions required to reach 95% of peak decoding accuracy. Results indicate functional and spectral (cvPCA-PR) dimensionality values converge on a similar low-dimensional PFC object representation

### PFC compresses high-dimensional IT representations

A low-dimensional representation in PFC could arise because it compresses high-dimensional inputs from visual cortex. Alternatively, it could inherit a representation from visual cortex that is already low-dimensional, due to compression earlier in the cortical hierarchy or to low-dimensional structure in the objects themselves. The PFC-compression hypothesis predicts that, for the same set of objects, PFC dimensionality should be much lower than that of its inputs from inferotemporal (IT) cortex. The inheritance hypothesis predicts dimensionality in both areas should be similar.

Although we did not record from visual cortex in this experiment, we used a well-established computational proxy for IT to estimate the dimensionality of its object representation. We used the pre-logit embeddings of the ResNet50 deep neural network, which has been shown to well approximate IT population responses [46,47]. This convolutional neural network (CNN) was trained to recognize objects, and developed internal representations that closely resemble IT neural responses. Two partially overlapping stimulus sets (set A,B) were used across the 18 sessions (set A was used in all sessions except one). Because ResNet50 embeddings are deterministic (i.e. there’s no trial-to-trial noise), per-session PR would be redundant. We therefore computed the mean participation ratio per stimulus set using standard (non-cross-validated) PCA on each set’s 40-object embedding matrix.

In PFC, variance was strongly concentrated in the first few principal components (Fig 6A, blue and red). In contrast, for the IT proxy, variance was much more distributed across components (Fig 6A, purple). Accordingly, the IT proxy had a mean PCA-PR dimensionality of 25.6 across the two stimulus sets used in our experiments (range 23.7–27.5; Fig 6B). Compared to PFC’s cvPCA-PR dimensionality values of ∼3–6 (this is significantly greater *p* = 3.8 × 10^−6^; one-sided Wilcoxon signed-rank test for both sample and delay) and represents an approximately 5–7 fold reduction in effective dimensionality from IT to PFC. This suggests that PFC does not inherit a low-dimensional object representation from IT or earlier processing, but instead that there is a substantial compression from IT to PFC.

**Fig 6.**
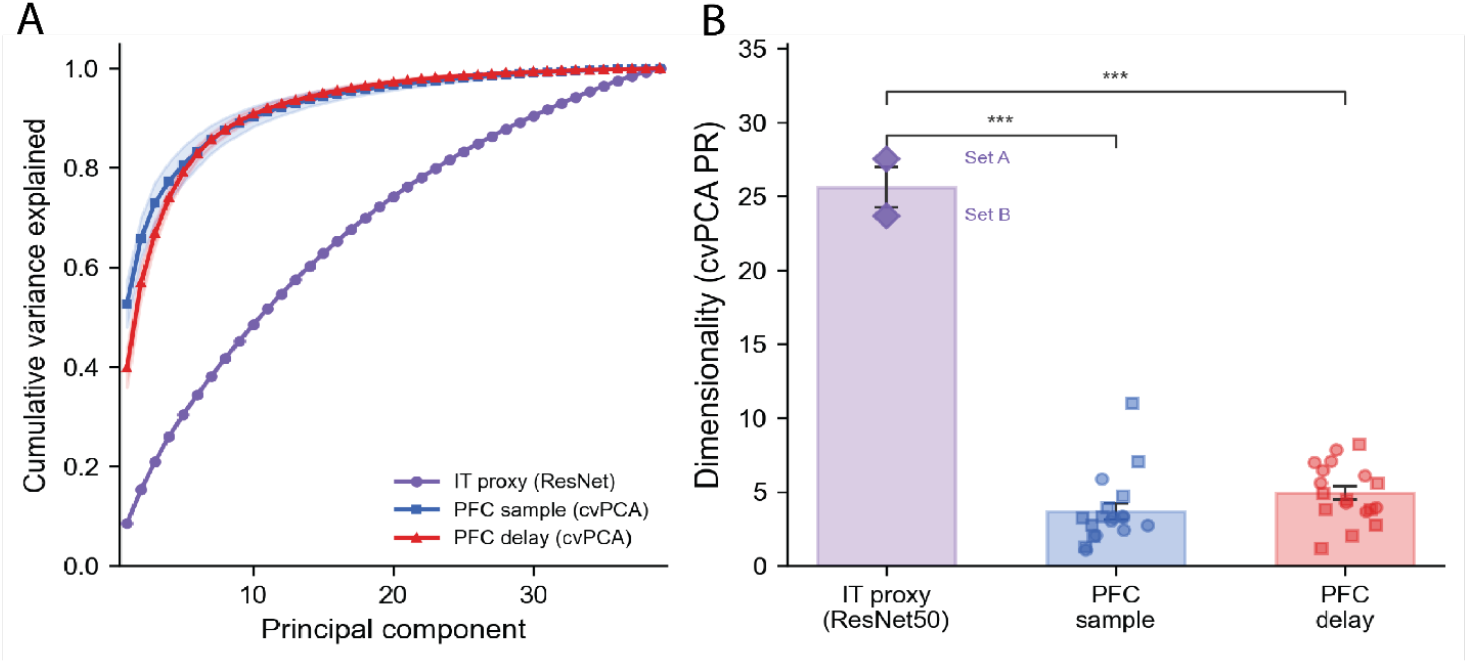
PFC object representations are lower-dimensional than a well-established inferotemporal cortex (IT) model. (**A**) Cumulative variance explained across principal components for ResNet50 pre-logit embeddings used as an IT proxy and for PFC sample and delay period activity. Variance in PFC is concentrated in the first few components while the IT proxy requires many more dimensions to explain the same fraction of variance. (**B**) Effective dimensionality (cvPCA-PR) for the IT proxy as well as PFC sample and delay epochs. Bars indicate mean ± SEM and circle and square points indicate session-wise values for each subject respectively. The IT proxy is significantly more high-dimensional than both PFC epochs. For the IT proxy bar, we plot each set’s PR as a diamond and we set the bar’s height to be the unweighted mean of the two. The SEM reflects variability between stimulus sets and is not directly comparable to the PFC whiskers, which are SEMs across recording sessions. Together these results support the hypothesis of PFC “compressing” higher-dimensional visual object representations into a functionally relevant low-dimensional code.

### From encoding to maintenance: dimensionality shifts and geometric preservation

To examine how dimensionality evolves with finer temporal resolution, we computed time-resolved cvPCA-PR in sliding 500 ms windows (100 ms steps) spanning the full trial. To exclude time points where there was no object information in the population activity — and thus the effective dimensionality was undefined — we only included times where cross-validated signal variance was significantly greater than its trial-shuffled control.

We found that dimensionality remained relatively stable over the course of the working memory delay period (Fig 7A). Following a small initial transient increase at sample onset for one subject (subject D), dimensionality remained relatively constant, with a very slight increase through the time course of the trial. To quantify the trend over the delay period, we fit a linear slope to each session’s curve of cvPCA-PR over time across the delay epoch and tested whether slopes differed from zero across sessions. Slopes showed positive trends for both subjects, but were only significant for one subject (Subject D: slope = +4.23 ± 1.01 dimensions/s, p = 0.008; Subject T: +0.94 ± 0.66, p = 0.20). These results suggest dimensionality exhibited at most a modest increase over the time course of the delay.

**Fig 7.**
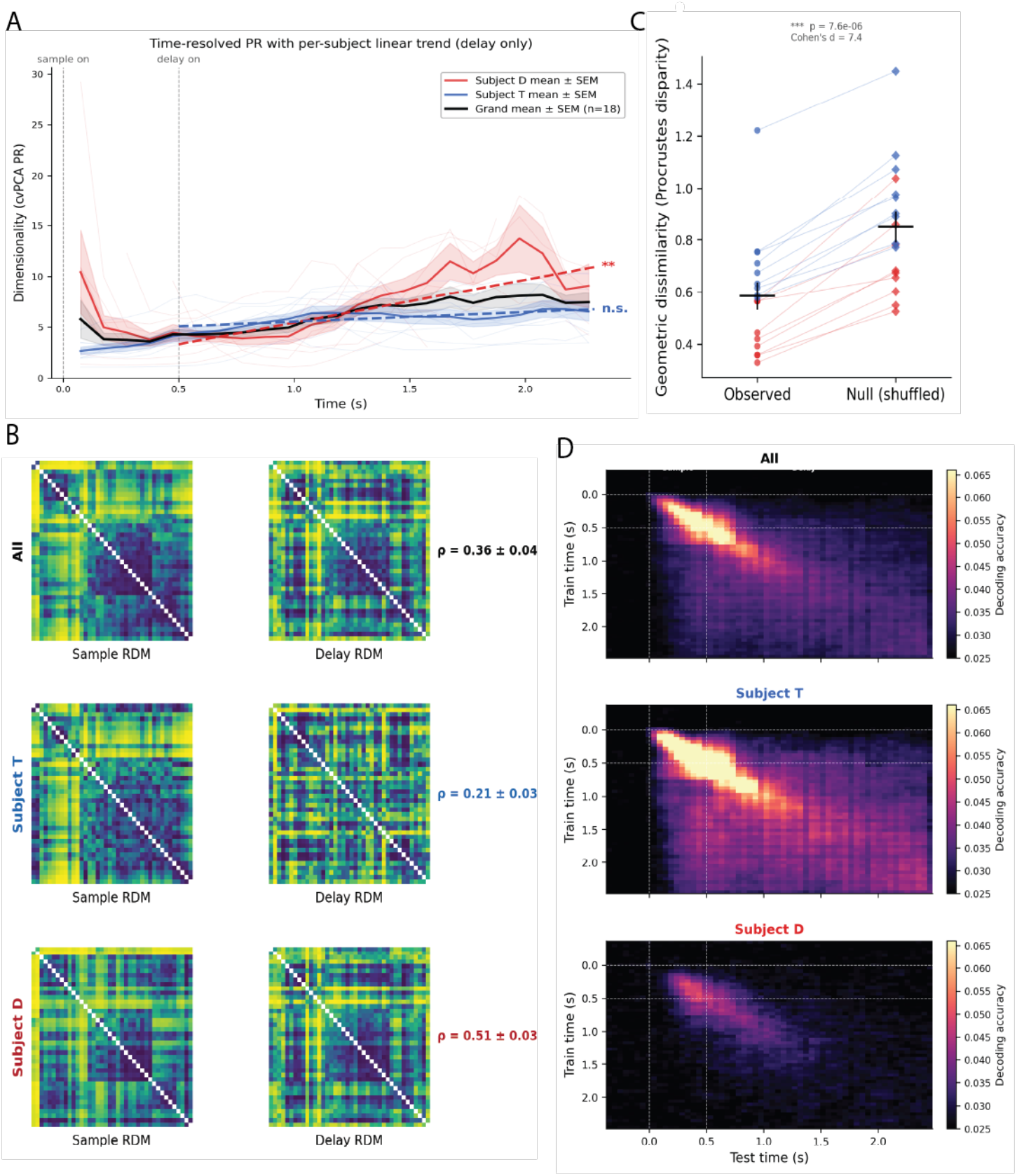
Object representations remain low-dimensional from encoding through memory maintenance. (**A**) Time-resolved dimensionality (cvPCA-PR) across the trial. Thin colored lines show individual sessions, colored by subject. Bold blue and red traces show subject mean ± SEM, and the thick black trace shows the grand mean ± SEM across all sessions. Dimensionality showed a modest increase over the course of the delay epoch (only significant for subject D). (**B**) Representational dissimilarity matrices (RDMs) showing dissimilarity between all pairs of objects, for both the sample and delay epochs, and for all sessions combined and separately for each subject. RDMs were significantly correlated across epochs, indicating PFC representational geometry for objects is somewhat preserved. (**C**) Procrustes disparity between sample and delay representations. Observed disparities were lower than shuffled null values, confirming that the geometry of object representations was preserved more than expected by chance. (**D**) Cross-temporal decoding matrices for 40-way object identity classification, shown for all sessions combined and separately for each subject. Color indicates decoding accuracy for training at one time point and testing at another. Cross-epoch generalization of decoding was weak, indicating that encoding and maintenance rely on partially distinct population subspaces.

The geometry of the PFC object representation was also largely consistent across time. To quantify this, we computed the relationship between population activity patterns for each object using representational similarity analysis (RSA) [48]. For each epoch, we computed the dissimilarity (Euclidean distance) in population activity space between condition-mean responses for all pairs of objects. This resulted in an object × object representational dissimilarity matrix (RDM) for each of the two epochs (Fig 7B). We then computed the Spearman rank correlation between these two matrices. This demonstrated that the relative geometric arrangement of objects in PFC population space was also largely preserved across the sample and delay epochs (cross-validated RSA: *ρ* = 0.363 ± 0.041, *p* = 7.6 × 10^−6^).

We used Procrustes analysis to confirm that this geometric preservation extends beyond pairwise distances to the overall spatial configuration of the 40 object centroids. Procrustes alignment measures how well one geometric configuration can be rotated and reflected to match another. The residual mismatch after optimal alignment is the Procrustes disparity, which is zero when two configurations have identical shapes. We found that the disparity between the PFC sample and delay epoch object representations (0.584 ± 0.049, mean ± SEM) was significantly lower than a label-shuffled null that broke the object-identity correspondence between epochs (0.849 ± 0.055; Wilcoxon, *p* = 7.6 × 10^−6^; mean Cohen’s *d* = 7.4; Fig 7C). The RSA and Procrustes analyses together indicate that the spatial arrangement of objects within PFC population activity is substantially preserved across working memory encoding and maintenance.

Although the dimensionality and representational geometry were relatively stable, the PFC neural code for objects did undergo a substantial reconfiguration between encoding and maintenance. To quantify this, we ran a cross-temporal decoding analysis, training and testing an SVM object classifier at all pairs of time points. The resulting cross-temporal matrix (Fig 7D) showed weak generalization of decoding across the sample and delay epochs, as has been previously observed [49–51]. This suggests encoding and maintenance rely on partially distinct population subspaces. Quantifying this subspace shift, the cross-validated leading principal angle between the sample and delay object subspaces (K=6, matched to the cvPCA PR) increased across ranks: 33.7° ± 2.9°, 44.9° ± 2.9°, 54.4° ± 2.7°, 65.1° ± 2.2°, 76.4° ± 1.0°, and 85.7° ± 0.4° (mean ± SEM, n = 18 sessions). Every rank was significantly above a within-epoch noise floor estimated from split-halves of the same epoch (one-sided Wilcoxon signed-rank, all p_Bonferroni < 10^−4^, 18/18 sessions positive at every rank). Taken together, these results are consistent with a rigid rotation of the PFC population activity which preserves the dimensionality and the representational geometry.

## Discussion

We measured the dimensionality of object representations in the prefrontal cortex during a working memory task. We found that a set of 40 objects could be captured in only ∼3–6 effective dimensions, far below the theoretical maximum. Multiple control analyses confirmed that this low dimensionality was not due to limited sampling of neurons or conditions. A comparison with a neural network proxy for inferotemporal cortex (IT) suggested that PFC compresses its IT inputs by ∼5–7 fold. Despite an overall rotation of the population activity patterns coding for objects between working memory encoding and maintenance, the dimensionality and representational geometry were preserved.

### Sampling and experimental design can influence dimensionality

Estimates of neural population dimensionality can depend to some degree on how they are measured. Dimensionality is bounded by the number of independent conditions tested, and can also be underestimated if the actual dimensionality is close to the number of conditions [20,37,42,52,53]. *C* conditions span at most *C*–1 directions in population activity state space, and as the true dimensionality approaches this ceiling, the smallest signal modes shrink toward the noise floor and are discarded as noise. In primary visual cortex (V1), it has been shown that estimated dimensionality increases essentially indefinitely with the number of visual images sampled [20] (although recently there have been alternative interpretations to these results [21]).

In contrast, we found that PFC exhibits clear saturation: The effective dimensionality plateaued well before reaching the limit imposed by the number of conditions we tested. This indicates that our tested conditions were sufficient to accurately measure PFC dimensionality. This suggests that PFC indeed represents objects within a fixed low-dimensional subspace.

The degree of compression is also likely to depend on task demands [54,55]. Recent work showed that object-response reinforcement learning resulted in higher-dimensional PFC activity than location-response learning [55]. Tasks that require a reduction to binary (i.e., one-dimensional) cognitive representations, tend to result in essentially one-dimensional representations in higher-level areas like PFC [9,56,57]. Hypothetically, a task requiring more detailed, fine-grained object discriminations might conceivably result in a higher-dimensional representation.

However, our commonly-used object delayed-match-to-sample task [1,31–34] requires basic-level object recognition [58–60] and maintenance in working memory. That is, it only required subjects to maintain sufficiently detailed object representations to discriminate between different object identities. We suggest this may represent a default level of PFC object representation in the absence of specific learning to generalize to broader categories or make finer discriminations. In this regime, we found a low-dimensional PFC object representation that is between these two extremes.

Dimensionality estimates can also depend to some degree on the analytical methods used to estimate them. We observed a small difference between spectral (cvPCA-PR; ∼3-6) and functional (classification-based; ∼6-10) estimates of PFC dimensionality. This might be due to intrinsic differences in the way these methods estimate dimensionality. Or it might be due to the fact that cvPCA explicitly corrects for noise, while the classification-based method does not. Nevertheless, we suggest that more important than the precise number of dimensions is the general idea, consistent across both methods, that PFC represents objects with only a small handful of dimensions.

### Dimensionality decreases across the cortical hierarchy

Though dimensionality estimates can vary widely across tasks, analytical methods, and brain areas, a consistent pattern has been observed across the cortical hierarchy. Early sensory areas often maintain high-dimensional representations of inputs from the periphery [19,20]. Work in visual cortex areas V4 [61] and IT [41,62] suggests that dimensionality is compressed through the cortical hierarchy [63], though it remains relatively high even at later stages of the ventral stream [38]. By contrast, in frontoparietal cognitive and motor areas, neural activity is well-captured by low-dimensional manifolds [8,13,15,17,64–66]. This suggests that task-relevant variables may be further compressed into a smaller latent space.

Consistent with this idea, for the same set of objects, our neural network proxy for IT showed ∼26 effective dimensions, while PFC data showed only ∼3–6. The extent to which dimensionality is reduced in a gradual, stepwise manner across the cortical hierarchy [63] or whether there is a sharper decrease in dimensionality between visual cortex and PFC, as suggested by some studies of categorization [56], is an open question for future work. Taken together, these results suggest that instead of preserving the detailed structure of sensory stimuli, PFC maintains a more compressed representation of object identity, which may be more suitable for working memory and decision making.

### Mechanisms of dimensionality reduction

Converging connectivity from the visual cortex to the PFC could explain the compression to low dimensionality. That is, PFC could receive a low-dimensional projection of visual cortex outputs. This might correspond to random convergent projections [67–70], analogous to reservoir computing models [71]. Alternatively, IT-PFC connectivity might selectively emphasize a small set of behaviorally relevant visual dimensions, as suggested by studies of categorization [56,57].

Another contribution to PFC dimensional compression could be recurrent connectivity within PFC itself. Work on low-rank recurrent neural networks shows that the dimensionality of population activity is constrained by the rank (dimensionality) of the structured component of recurrent connectivity [72]. Low-rank connectivity also has biological relevance—it can be easily learned via Hebbian plasticity [73]. Within such low-rank networks, recurrent dynamics can implement attractors, neural state-space regions toward which population activity evolves and stabilizes. Attractor networks have been frequently proposed as models for working memory in PFC [74–76], as well as in other domains [77,78]. In these frameworks, the effective “memory” variable typically lives on a low-dimensional manifold, with the activity’s variability concentrated along these dimensions. Disentangling the relative contributions of low-rank readout vs recurrent attractors will likely require targeted perturbations and circuit-level modeling.

### Implications for prefrontal coding and working memory

Previous attractor models of working memory have typically been limited to 1D attractors to explain working memory of variables that are intrinsically continuous and one-dimensional, such as spatial location (polar angle) [74,76] and color (hue) [69]. Other models have used discrete attractors to maintain simplified low-dimensional tokenized inputs [79–81]. Our results suggest that low-dimensional attractor models may also be more generally applicable to working memory for objects, which are intrinsically very complex and high-dimensional [38–40]. A hallmark of attractor dynamics is a transient period of higher-dimensional activity immediately following stimulus onset, followed by convergence onto a lower-dimensional manifold as the attractor state is reached. This is consistent with the temporal pattern of dimensionality we observed in one subject (subject D), providing some evidence to support attractor dynamics for objects in PFC.

An open question is whether the low-dimensional object representation we observed is better described as a set of discrete attractor states [79–81] or as a continuous attractor manifold [69,74,76,82]. These two options have different implications. In the discrete attractor case, each encoded object might be slotted into one of multiple separate stable states in PFC activity [83]. Such architectures are highly robust to noise. But this robustness would come at the cost of discretizing similarity between different objects and requiring a separate attractor state for every new object encoded in memory. On the other hand, in a continuous attractor, objects are represented as positions along a shared low-dimensional manifold. Although continuous attractors may be more susceptible to drift and interference between memory states [7,74], they could preserve visual or behavioral similarity between objects and support generalization to novel stimuli. Distinguishing between these possibilities will be critical to a complete computational understanding of object working memory.

Our results may also, somewhat surprisingly, help explain the high-dimensional coding implied by nonlinear mixed selectivity [23,26,27]. Many studies have shown that prefrontal neurons respond to idiosyncratic combinations of sensory stimuli, contexts, and actions [10,22,24,25,84]. This “nonlinear mixed selectivity” results in a high-dimensional combinatorial code that permits downstream areas to flexibly read out virtually any arbitrary function of information coded in PFC [22,27,28]. But lateral PFC is a nexus of essentially all sensory modalities [85,86], action selection for a wide variety of effectors [87–89], and much of the cognitive and control-related processing in between [90,91]. If PFC were to faithfully maintain the full dimensionality of each of these variables, this would likely entail a combinatorial explosion that would far exceed the number of available neurons. Our results suggest that PFC may adopt a strategy by which it compresses the dimensionality of each individual variable it encodes, in order to make their combination into a high-dimensional mixed code tractable.

## Conclusion

Our results suggest object representations are compressed into a low-dimensional manifold in PFC. This compression may reflect prefrontal attractor dynamics, and might facilitate interaction of object representations with other variables and cognitive control. These results illuminate an important aspect of prefrontal coding and provide key constraints for models of working memory.

## Methods

### Non-human Primate (NHP) subjects

Two adult macaque monkeys (*Macaca mulatta*), monkey T (male, 17 years, 13 kg) and monkey D (male, 8 years, 10 kg) served as subjects in this study. The NHPs were pair-housed, on a 12-hr day/night cycle, and in a temperature-controlled environment (80° F). They were enriched with a diverse range of toys, puzzles, novel foods, and social interaction with other NHPs and humans. All procedures followed the guidelines of the National Institutes of Health and the Massachusetts Institute of Technology Committee on Animal Care (protocol 2501000771).

### Behavioral Task Paradigm

We ran a delayed match-to-sample object working memory task. Following a brief fixation period (500 ms), a sample object was presented for 500 ms. The NHP subjects were then required to maintain the sample object in working memory over a 2 s blank delay period. Finally, two test objects — one matching the sample and a randomly selected non-matching object — were shown 7.5° of visual angle from fixation, at random diametrically-opposed angles. Subjects were allowed to freely view both test objects before indicating their choice by fixating on it for at least one second. Correct choice of the matching test object was reinforced with a juice reward. Subjects performed very well on this task (mean behavioral performance across sessions was 99.6% for subject T, 86.8% for subject D).

Samples and test objects were selected from a pool of 40 full-color natural object images isolated from their background. Images were licensed from a royalty-free commercial photo database (Hemera Photo-Objects, Vol. 1 & 3). All sessions except one used the same set of 40 object images (Fig 1C); a partially overlapping set of 40 objects was used in the remaining session (Supplementary Fig S1). Object images were resized so they all had approximately the same size (5° radial distance from center to farthest extent of object).

### Electrophysiological data collection

All data was recorded from four chronic 64-electrode microelectrode arrays (256 electrodes total) implanted bilaterally in dorsolateral PFC (dlPFC) and ventrolateral PFC (vlPFC). We used iridium-oxide “Utah” arrays (Blackrock Microsystems, Salt Lake City, UT) with an 8 x 8 electrode layout, 400 µm spacing, and 1 mm length. Electrodes in each hemisphere were grounded and referenced to a separate subdural reference wire. Spiking activity was amplified, filtered (250–5,000 Hz), digitized, manually thresholded to extract spike waveforms, and streamed to disk at 30 kHz (Cerebus, Blackrock Microsystems). Spike waveforms were sorted online into putative units using multiple time-amplitude windows.

### Data selection and preprocessing

Eight electrodes with identical signals, due to apparent cross-talk, were removed from all sessions of subject T. Occasional bouts of noise induced simultaneous spike-threshold crossings in the raw data across most or all channels on an array. We identified these as any time points with threshold crossings on ≥3 electrodes on an array at the same exact 30 kHz (0.03 ms) timestamp, and removed them from all channels on that array. Units with overall spike rates < 0.01 spikes/s were excluded from analysis. All analyses only include correctly completed trials. Prior to all analyses, except for fitting the Poisson-based generative population model, each unit’s spike rates were z-scored based on the mean and SD across all trials during the 500 ms fixation period baseline.

### Cross-validated PCA (cvPCA)

We used cross-validated PCA [20] (cvPCA) to isolate reliable stimulus-related signals from any noise in estimating condition-mean neural activity. We split trials for each condition (sample object) into two equally sized subsets: *A* and *B*. We computed condition means separately for each split-half subset, and mean-centered these values across conditions, resulting in two [conditions × neurons] condition-centroid matrices: ***M***_*A*_ and ***M***_*B*_. We performed Singular Value Decomposition (SVD) on ***M***_*A*_ to get a set of *p* eigenvectors ***v***_*A,k*_. Then we projected the split-half-*B* condition means ***M***_*B*_ onto each split-half-*A* principal direction ***v***_*A,k*_ and computed the *k*^*th*^ cross-validated eigenvalue as:

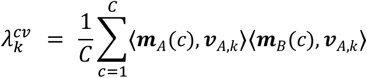

where ***m***_*p*_*(c)* and ***m***_*B*_*(c)* are the *c*-th rows of ***M***_*A*_ and ***M***_*B*_ (i.e., the centroid for condition *c* in each split), and *C* = 40 is the number of conditions. We repeated this procedure for 50 independent random trial splits. This yielded a [50 x (40 – 1)] matrix of cross-validated eigenvalues.

Because the noise in trial subsets *A* and *B* is independent, the expected value of the product of the noise across subsets, *E*[*noise*_*A*_ ⋅ *noise*_*B*_] is 0. Thus, the cvPCA method results in an unbiased estimate of the signal variances [20].

### Participation Ratio

The participation ratio [42] (PR) quantifies the effective dimensionality of a given distribution of variance across principal components. Given a set of eigenvalues *λ*_*1*_ ≥ *λ*_*2*_ ≥ ⋯ ≥ *λ*_*p*_ from a covariance matrix, the PR is computed as:

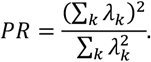

The numerator is the square of the total variance. The denominator is a measure of how concentrated that variance is. Their ratio is a continuous measure of how many dimensions carry substantial variance. PR equals 1 when all variance is concentrated in a single dimension, and equals the maximal number of dimensions when variance is uniformly distributed. We computed the participation ratio from the mean cross-validated eigenvalues returned by cvPCA, averaged across all 50 trial splits. This constituted a pipeline for computing the signal-only dimensionality, which we refer to as cvPCA-PR.

### Condition-permutation null

To test whether the observed cvPCA-PR dimensionality values exceed the level expected from estimation noise alone, we constructed a condition-permutation null. Within each of the 50 trial splits, the condition labels of the half-*B* condition-mean centroids were randomly permuted before computing cross-validated eigenvalues. Because the permutation removes the correspondence between the object identities *c* in half *A* and the permuted objects *c*′ in half *B*, the projections (***M***_*A,c*_ ⋅ ***v***_*k*_) and (***M***_*B,c*′_ ⋅ ***v***_*k*_) are uncorrelated. Thus, all eigenvalues have expected values of 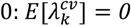 for all *k*. The resulting null eigenvalues fluctuate symmetrically around zero and the PR collapses to a small positive value reflecting estimation noise alone.

Note that the signal cvPCA-PR and the null PR values are computed at different levels of aggregation. The signal PR is computed from the mean eigenspectrum averaged across all 50 splits. Each null PR is computed from a single permuted eigenvalue array from one split, preserving the full sampling variability of the null. We generated approximately 20 permutations per split x 50 splits = 1000 null PR values per session. Per session p-values were computed as the fraction of these 1000 null PRs that equalled or exceeded the observed signal PR. To assess population-level significance, we performed a one-sided Wilcoxon signed-rank test on the difference 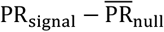 (where 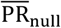 is the mean of the 1,000 null PR values) across all 36 session–epoch pairs (18 sessions × 2 epochs).

### Dimensionality as a function of number of conditions

To examine how estimated dimensionality varied with the number of conditions tested, and whether it had saturated at the 40 conditions we tested, we computed cvPCA-PR on subsets of sample object conditions. We randomly subsampled *C*′ = *3,5,8,10,15,20,25,30,35,40* objects (5 subsamples per *C*′, 25 cvPCA splits per subsample). To characterize this function, and extrapolate it to an infinite number of conditions, we fit a saturating exponential to this data for each subject and epoch:

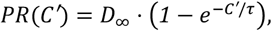

where *D*_∞_ is the asymptotic dimensionality and *τ* the characteristic number of objects at which saturation occurs. The asymptote *D*_∞_ estimates the dimensionality in the limit *C* → ∞. Model fitting used nonlinear least-squares (`scipy.optimize.curve_fit`). This model fit the data well (all *R*^*2*^ > *0*.*97*), better than other functional forms we tested. 95% confidence intervals on *D*_∞_ were obtained by bootstrap: sessions were resampled with replacement 1000 times, the cross-session mean curve was reconstructed on each bootstrap and the 2.5-97.5 percentile of the resulting *D*_∞_ distribution was taken as the 95% confidence interval.

### Dimensionality as a function of neuronal population size

To examine how estimated dimensionality varied with the number of neurons sampled, and whether it had saturated with the population sizes we recorded of *N*∼100 neurons, we computed cvPCA-PR on subsets of neurons. For each session and each epoch, we randomly sampled *N*′ = *10* to *N* neurons at 20 geometrically-spaced points (20 subsamples per *N*′, 50 cvPCA splits) and measured the cvPCA-PR at each *N*′. At each population size, we computed a cross-session weighted mean of PR, weighting each session by its recorded population size. The SEM was the weighted population standard deviation divided by the square root of the number of contributing sessions. Because the maximum recorded population differed across sessions, the common range over which all sessions contributed extended only to the smallest per-session population within each subject (Subject T: 35 neurons, Subject D: 118 neurons).

A saturating exponential was fit to this function:

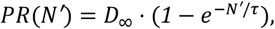

The asymptote *D*_∞_ estimates the dimensionality in the limit *N* → ∞. This model generally fit the data well (all *R*^*2*^ > *0*.*90*, except subject T sample epoch: *R*^*2*^ = *0*.*28*, which was essentially flat across *N*). These fits were better than other functional forms tested. 95% confidence intervals on *D*_∞_ were obtained by the same bootstrap procedure described above for the conditions saturation analysis.

### Generative population model

To determine whether dimensionality might increase for population sizes beyond what we sampled in our data, we used Monte Carlo methods to simulate synthetic population data with a generative model fitted to the covariance and rate statistics of the real data.

#### Data selection for fitting generative model

To ensure that our generative model was driven by high signal-to-noise ratio data, we pooled neurons across 8 sessions in subject T (we removed one session that had a different object set, see Supplementary information), whose data carried stronger object information (higher decoding accuracy). We used an iterative pruning process based on the decoding accuracy of the aggregated population to find the most informative neurons. For each epoch, we computed the condition-mean spike rates for each neuron separately for a training and testing data subset (5-fold cross-validation), and used these as features for a linear SVM classifier. In each iteration, we computed each neuron’s classification “importance” as the L2 norm of its SVM weight vector, averaged across cross-validation folds. The bottom 10% of neurons ranked by this importance score were then removed, and this procedure was repeated until cross-validated decoding accuracy fell below 95% of its peak value. When this stopping criterion was reached, we retained all neurons included in the previous (above-criterion) iteration. This procedure yielded a pseudopopulation of 139 informative neurons. The 40-way classification accuracy of these 139 neurons was 73.5% for decoding cross-validated condition centroids and 37% for decoding single trials (2.5% chance).

#### Zero-inflated poisson (ZIP) model for spike rate statistics

To model the condition-specific spike rate of each neuron, we fit the observed responses with a zero-inflated Poisson distribution [45,92]. This is conceptually similar to other “overdispersed” models, which have been used previously to successfully model neural activity that is more variable than predicted by a standard Poisson distribution [93,94]. This simple model builds on the standard Poisson distribution by incorporating an additional “zero-inflation” term that sets a rate of 0 with some probability. Thus, it is capable of capturing the varying degrees of response sparsity observed across neurons [95,96].

The spike count *y*_*i,c,t*_ for each neuron *i*, sample object condition *c*, and trial *t* is:

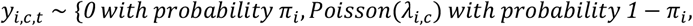

where *π*_*i*_ ∈ [*0,1*] is a neuron-specific, but condition-invariant, zero-inflation probability shared across objects. Parameters were estimated by coordinate-descent maximum likelihood with L1 regularization on *logλ*_*i,c*_ (regularization strength *α* selected by 5-fold cross-validation, resulting in selected 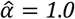).

Standard errors of the parameter estimates were obtained by Monte Carlo resampling (200 replicates): on each replicate, a synthetic dataset of *T*=23 trials per neuron-object pair (the minimum number of trials across sessions) was generated from the fitted parameters of the ZIP model and the full estimation procedure was rerun. The standard error of each parameter was computed as the standard deviation of its Monte Carlo distribution across replicates. These errors are used below to generate shrinkage estimates of the condition means.

#### James-Stein shrinkage of condition means

To reduce estimation noise in the rate profiles, we applied James–Stein shrinkage [97] on the ZIP-fitted values of 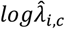. For each neuron, this procedure shrinks the object-specific rate estimates toward that neuron’s own grand-mean rate across objects, yielding a more uniform but lower-variance tuning profile. The degree of shrinkage is controlled by the ratio of the mean Monte Carlo variance (estimation uncertainty) to the total sum of squares of the tuning profile (signal spread). Neurons whose object-specific rate modulation is small relative to their estimation noise are shrunk more heavily. For neuron *i* and sample object condition *c*, let the estimated Poisson log-rate be 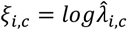, its mean across object conditions be 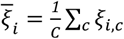 and its sum of squares across conditions be 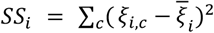. The shrinkage coefficient is 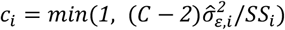), where 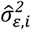 is the mean Monte Carlo variance (as described in the previous section). The shrunk estimate for each neuron *i* and object condition *c* is: 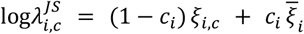.

#### Between-condition covariance estimation and multivariate Gaussian density fit

To generate novel simulated neurons beyond our recorded ones, we estimated the between-condition covariance matrix by fitting a multivariate Gaussian to the James-Stein shrunk rate profiles. Let 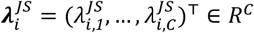 denote the vector of fitted and shrunken Poisson rates for each condition, for neuron *i*. The sample mean and covariance across the *N* = *139* real neurons are

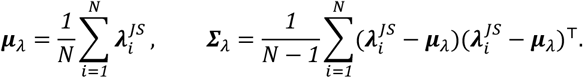

To generate a novel simulated neuron *j*, we randomly sampled rates from the resulting multivariate Gaussian distribution:

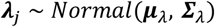

clipping ***λ***_*j*_ element-wise to a minimum of *0*.*01* to ensure valid Poisson rates.

A separate univariate Gaussian was fit to the zero-inflation probabilities 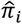 in logit space, matching the parameterization used in model estimation: 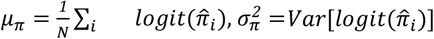. For each novel simulated neuron *j*, we also randomly sampled zero-766 inflation probabilities from this univariate Gaussian distribution:

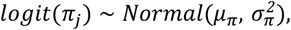

and apply the inverse logit to obtain *π*_*j*_ ∈ [*0,1*]. Per-trial spike counts were then generated from the sampled ***λ***_*j*_ and *π*_*j*_ based on the ZIP model above.

We used this generative model to sample synthetic neuronal populations of varying sizes up to 3,000 neurons. We ran the same cvPCA-PR and offset-exponential fit analytical pipeline on these that we used for the real data.

#### High-dimensional generative population model

To confirm that our cvPCA-PR analysis is able to successfully recover high-dimensional structure when it is present, we simulated a high-dimensional population using our generative model. We aimed to eliminate the between-condition covariance structure underlying the observed low dimensionality, while preserving the observed within-condition variances. Thus, we simply ablated (set to 0) all off-diagonal entries in the estimated between-condition covariance matrix ***Σ***_*λ*_, resulting in a diagonal covariance matrix ***Σ***^(*diag*)^_*λ*_. We then generated synthetic neuronal populations of varying sizes, as above, but instead sampling from the diagonalized multivariate Gaussian distribution: ***λ*** ^(*diag*)^_*j*_ ∼ *Normal*(***μ***_*λ*_, ***Σ***^(*diag*)^_*λ*_).

### Classification-based dimensionality (dim_class_)

To estimate dimensionality using an alternative, information-based method, we computed the object decoding accuracy on data projected onto varying numbers of its own principal components. For each session and epoch and cross-validation fold (stratified 5-fold cross-validation repeated 10 times), PCA was computed on the training trial subset only, in two steps. First we computed the condition-averaged responses from the training trials. We applied mean centering and then we used SVD to obtain *K* = *C* − *1* = *39* orthogonal axes in population activity space. Then, we ordered these axes by their cross-validated eigenvalue magnitude |*λ*_*cv*_| in descending order, using the same split-half procedure as the main cvPCA analysis. Because both these steps were computed on the condition-averaged responses (centroid) from the training set, the projection basis was estimated completely independently from the held-out test data.

For *k* = *1,2*, …,*39* dimensions, both training and test data were projected onto the top *k* cvPCA-ordered dimensions, the resulting features were standardized, and classification accuracy was computed using a linear SVM (LinearSVC, *C*_*reg*_ = *1*.*0*, balanced class weights). The resulting curves of decoding accuracy as a function of number of dimensions *k* were normalized to the range [0,100]% (zero to max) for each session, and averaged across sessions. The final information-based dimensionality estimate, *dim*_*class*_, was defined as the smallest number of PCA dimensions at which mean normalized decoding accuracy reached 95% of its maximal value across all *k*.

### Estimating inferotemporal cortex dimensionality via neural network model

To estimate the dimensionality of object representations in inferotemporal (IT) cortex without actual IT recordings, we used a well-validated computational proxy for IT. Previous studies [46,47] have shown that deep convolutional neural networks (CNNs) trained to recognize objects develop internal representations that closely resemble those found in IT, and they are capable of effectively predicting individual IT multi-unit responses. We used the activations of the penultimate (prelogit) layer of the 50-layer CNN known as ResNet50 [98]. These ResNet50 activations were generated for the same sets of objects used in our actual experiments, and standard PCA was used to estimate their dimensionality (non cross validated since the CNN object embeddings are deterministic and thus, contain no trial-to-trial noise).

### Sample vs Delay Epoch Comparisons

To compare dimensionality across the sample and delay epochs, we ran a paired Wilcoxon signed-rank test on their cvPCA-PR values across the 18 sessions. Additionally, the representational geometry comparison analyses test qualitatively different hypotheses (distance preservation, shape preservation, decoding stability).

#### Time-resolved cvPCA-PR with permutation thresholding

In order to track how dimensionality evolves more continuously over the time course of trials, we computed cvPCA-PR in sliding temporal windows (500 ms width, 100 ms step) spanning the full trial period. Since effective dimensionality is not well-defined at time points where there is little to no signal in the data, we excluded non-informative windows from this analysis. Within each window, we assessed whether the total cross-validated signal variance was significantly above chance using a within-split permutation test. For each of 30 split-half repetitions, 10 null samples were generated by shuffling condition labels within the split, yielding 30×10 = 300 null eigenspectra per window. The total null signal variance was computed for each null sample, and a one-sided *p*-value was obtained as the fraction of null signal variances ≥ the real signal variance. Windows where the total signal variance did not significantly exceed the null (*p* < 0.05) were excluded from the cross-session summary statistics. Cross-subject grand-mean traces were displayed only at time points where ≥ 5 sessions contributed significant estimates.

#### cvPCA PR slope test

To quantify whether there was an overall increasing or decreasing trend of dimensionality across the delay period, we fit it with a simple linear function. For each session *j* we restricted the analysis to the delay window *t* ∈ [*0*.*5, 2*.*3*] s and used ordinary least squares to fit:

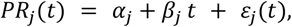

on the subset of windows with non-missing *PR*_*j*_(*t*) (requiring at least four windows per session). The session-specific slope *β*_*j*_ summarises the delay-period change in dimensionality. Per-subject group statistics were obtained by aggregating {*β*_*j*_}_*j*∈*subject*_ and reporting the mean ± SEM across sessions; statistical significance vs. the null hypothesis *β* = *0* was assessed with the two-sided Wilcoxon signed-rank test on per-session slopes.

The displayed regression line for each subject uses the within-subject means of session intercepts and slopes:

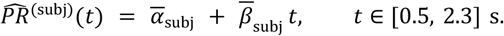

which is the proper visualization of the group fit and avoids conflating within- and between-session variance that would arise from fitting to the across-session mean curve. The grand-mean trend is computed analogously by pooling sessions across subjects.

#### Cross-temporal SVM decoding matrix

We used cross-temporal decoding to quantify how consistent the PFC population code for objects was across time within trials. For all pairs of time windows, a linear SVM (C=1.0, balanced class weights) was trained at one time point and tested at the other, using 5-fold stratified cross-validation repeated 2 times. Within each fold, each neuron’s firing rate was z-scored using the training fold statistics, and the same transformation was applied to the test fold.

#### Principal angles between sample and delay subspaces

Per session, we computed the principal angles between the top-6 PC bases of the per-epoch centroid matrices, estimated from disjoint trial halves (50 stratified per-object splits) to avoid trial-noise contamination; a within-epoch split-half procedure served as a finite-trial noise floor. Per rank, a paired one-sided Wilcoxon signed-rank test (Bonferroni-corrected across 6 ranks) compared the cross-validated sample-to-delay angle against this null across the 18 sessions.

#### Procrustes Alignment

To test whether the geometric arrangement of objects is preserved from sample to delay epochs, we used orthogonal Procrustes analysis. For each session, the condition-mean activity (centroid) from both epochs, ***M***_*sample*_and ***M***_*delay*_, was projected into a shared PCA space (fit on concatenated matrices and retaining enough dimensions to explain >99% of their combined variance). Each resulting projected centroid geometry was then scaled to unit Frobenius norm, so that their comparison would reflect shape differences, rather than overall magnitude. Orthogonal Procrustes analysis was then used to fit the rotation matrix ***R*** that minimizes the “Prorustes disparity”: 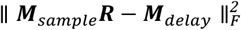. Significance was calculated by permuting the object labels of the delay epoch centroids.

#### Cross-validated Representational Similarity Analysis (RSA)

To further compare the representational similarity between sample and delay epochs, we computed and compared representational dissimilarity matrices (RDMs) for each epoch. Cross-validated Euclidean distances were computed using split-half condition-averaged population activity [99]. For each pair (*i,j*) of conditions, the cross-validated distance was the product of the half-*A* and half-*B* Euclidean distances,

*d*^*2*^_*cv*_(*i, j*) = ‖***m***_*A*_(*i*) − ***m***_*A-*_(*j*)‖ · ‖***m***_*B*_(*i*) − ***m***_*B*_(*j*)‖, where ***m***_*A*_(*i*) and ***m***_*B*_(*i*) are the condition centroids (rows of ***M***_*A*_ and ***M***_*B*_) for condition *i* within trial splits *A* and *B*, respectively. This resulted in an estimate of squared-distance that avoids the positive bias of distance of non-cross-validated distance estimates on noisy condition-averaged activity [99]. The Spearman rank correlation between the upper triangles of sample and delay RDMs was computed for each of 50 random trial splits and averaged.

### Computation

All analyses were performed in Python 3.13, using the NumPy 2.4, SciPy 1.17, scikit-learn 1.8, matplotlib 3.10, Pandas 3.0, statsmodels 0.14, PyTorch 2.12, and Spynal [100] libraries.

## Acknowledgements

We thank Jordan G. DeFarias for technical assistance in data collection. We thank Haim Sompolinsky, Nacho Castillejo, Jefferson Roy, and the rest of the Miller Lab for helpful comments and suggestions.

## Data availability statement

The data reported in this paper will be shared by the corresponding author upon reasonable request.

